# Drug-repurposing screen identifies thiostrepton as a novel regulator of the tumor suppressor DAB2IP

**DOI:** 10.1101/2024.07.12.603185

**Authors:** Rossella De Florian Fania, Serena Maiocchi, Raffaella Klima, Valeria Pellegrini, Sabrina Ghetti, Davide Selvestrel, Luca L. Fava, Luca Braga, Licio Collavin

## Abstract

The tumor suppressor DAB2IP, a RasGAP and cytoplasmic adaptor protein, modulates signal transduction in response to several extracellular stimuli, negatively regulating multiple oncogenic pathways. Accordingly, the loss of DAB2IP in tumor cells fosters metastasis and enhances chemo- and radio-resistance. DAB2IP is rarely mutated in cancer but is frequently downregulated or inactivated by multiple mechanisms. Solid experimental evidence show that DAB2IP reactivation can reduce cancer aggressiveness in tumors driven by multiple different oncogenic mutations, making this protein an interesting target for anti-cancer therapy. Based on these premises, we screened a library of FDA-approved drugs to search for molecules that can increase DAB2IP protein levels. We exploited CRISPR/Cas9 gene editing to generate two prostate cancer cell models in which endogenous DAB2IP is fused to HiBiT, a peptide tag that enables luminescence-based detection of protein levels in a sensitive and quantitative manner. Using this approach, we identified drugs able to increase DAB2IP levels. We focus our attention on thiostrepton, a natural cyclic oligopeptide antibiotic that has been reported to inhibit survival of various cancer cell lines. Functional experiments revealed that the cancer inhibitory effect of thiostrepton is reduced in the absence of DAB2IP, suggesting that the observed upregulation contributes to its action. These findings encourage the further development of thiostrepton for the treatment of solid cancers, and unveil a novel molecular mechanism underlying its anti-tumoral action.

## Introduction

Prostate cancer is one of the most common cancer in men; with an age-related incidence, it is the second leading cause of cancer death in men in the United States [1]. Current approaches are often efficient, but there are patients who do not completely recover from the disease. In fact, some patients develop resistance to chemo- and radiotherapy, rendering the treatment inefficient or even detrimental. Developing novel strategies to counteract the intrinsic causes of such resistance could significantly improve future therapies.

DAB2IP (Disabled-2 Interacting Protein) is a cytoplasmic Ras GTPase-activating (GAP) and an adaptor protein involved in signal transduction by multiple inflammatory cytokines and growth factors, negatively modulating key oncogenic pathways, such as TNF/NF-kB, WNT/β-catenin, PI3K/AKT, and Androgen Receptors [2, 3]. DAB2IP is also known as ASK1-interacting protein (AIP1), for its role in inducing the release of ASK1 from the inhibitory binding of 14-3-3 in response to TNF-α, favoring consequent activation of the pro-apoptotic ASK1-JNK pathway [4]. The loss of DAB2IP functions represents an advantage for cancer initiation and progression and accordingly it is frequently downregulated in various human malignancies [5, 6]. Multiple mechanisms of DAB2IP inactivation have been identified in cancer cells, suggesting that tumor cells take advantage from the inhibition of a single protein that negatively modulates various oncogenic signals. Intriguingly, DAB2IP is rarely inactivated by deletion or mutation, but it is preferentially inhibited at transcriptional and post-transcriptional level via different mechanisms [5, 6]. Various studies demonstrated that restoring DAB2IP expression in DAB2IP depleted cancer cells reverts metastatic behavior and drug-resistance both in vitro and in vivo, in multiple tumor types. Based on these evidences, DAB2IP is a strong candidate for development of therapeutics aimed to increase its protein levels in cancer cells, restoring its onco-suppressive role. In fact, even a moderate increase in DAB2IP levels may be able to limit cancer aggressiveness, as we could demonstrate by targeting some known mechanisms of DAB2IP inactivation [7–9].

Building upon the aforementioned concepts, in the present study we tested 1280 Food and Drug Administration (FDA)-approved drugs for their potential to increase DAB2IP levels in prostate cancer cell lines.

To monitor the amount and dynamics of the endogenous DAB2IP protein through a sensitive, quantitative, and high-throughput assay, we labeled endogenous DAB2IP with the luminescence-based tag HiBiT [10].

Using this approach, we uncovered candidate molecules able to modulate DAB2IP levels in cancer cells. We focus on thiostrepton, and provide evidence that DAB2IP upregulation contributes to its ability to counteract aggressiveness of cancer cells.

## Results

### Endogenous DAB2IP tagging with HiBiT

To discover drugs that can modulate DAB2IP, we tagged endogenous DAB2IP with the HiBiT peptide for quantitative, luminescence based detection of protein levels under physiological conditions. HiBiT is an 11 amino acid peptide that can be complemented with a larger subunit, LgBiT, to reconstitute a functional NanoLuc enzyme [10]. Since strong evidence indicate that DAB2IP reactivation in prostatic cancer counteracts metastasis and chemoresistance [11, 12], we chose to edit PC3 and DU145 human prostate cancer cell lines; the two lines differ for DAB2IP expression levels, p53 status, and overall aggressiveness, rendering them proper for exploring various mechanisms of DAB2IP modulation in cancer.

The human DAB2IP gene is complex; it has multiple transcriptional start sites, encoding at least three N-terminal variants of the protein; moreover, alternative splicing generates two possible C-termini (Fig. S1A). Aiming for a C-terminal fusion of the tag, we analyzed expression of the alternatively spliced variants and confirmed that both are co-expressed in several normal and transformed cell lines (Fig. S1B); therefore, to tag all possible DAB2IP isoforms, we placed the HiBiT peptide followed by a stop codon immediately upstream to the last donor splice site, so that the same C-terminally tagged DAB2IP protein is translated regardless of alternative splicing of the last intron (Fig. S1C).

To verify that the HiBiT peptide in such position would be functional, we cloned the corresponding construct in a mammalian expression vector and confirmed that DAB2IP could be readily detected by luminescence in transfected cell lysates (Fig. S1C-D).

We tagged endogenous DAB2IP with the HiBiT peptide in PC3 and DU145 using a CRISPR-Cas9 protocol previously described [13]. Bulk nucleofected cells were screened first by luciferase assay (Fig. S2B) and then by PCR on genomic DNA (Fig. S2C), confirming the successful integration of the HiBiT-tag into the target region. PC3 and DU145 cells positive for HiBiT insertion were selected as single clones with homozygous DAB2IP tagging. Responsiveness of the model was evaluated by siRNA-mediated silencing of DAB2IP, and by transfection of DAB2IP-targeting miR-149-3p [7]. The reduction of luminescence was in line with that observed by western blot, confirming that the system is applicable to measure variations in endogenous DAB2IP protein levels (Fig. S2D).

### High-throughput luminescence and fluorescence-based screen identifies molecules able to modulate DAB2IP expression levels

To screen for molecules that may increase DAB2IP protein levels, we used the more aggressive PC3 cells, that display lower basal expression of DAB2IP with respect to DU145. To normalize luciferase activity for variability in cell numbers, due to uneven cell seeding or toxicity promoted by the compounds, we quantified viable cells using a fluorescence-based assay (CellTiter-Fluor, Promega) that is compatible with subsequent luciferase quantification. We thus expressed endogenous DAB2IP levels as the ratio of Chemiluminence over Fluorescence intensity (Lum/Fluo) (Fig. S2 E-G).

We screened a library containing 1280 compounds most of which FDA- and EMA-approved (https://www.prestwickchemical.com). Each compound was added at 10 μM final concentration to the culture medium in 384-well plates as schematized in Figure 1A. For each drug, the Lum/Fluo ratio was normalized to control DMSO-treated cells and expressed as Z-score. Given the tumor suppressive activity of DAB2IP, we considered as potential hits also compounds that increased the Lum/Fluo ratio with an inhibitory effect on cell viability (Fig. 1B). According to these criteria, we selected 6 positive and 3 negative regulators that are summarized in Fig. 1B. After a secondary screen, 6 molecules confirmed their action (Fig. 1C). To corroborate these results we monitored the effects of the compounds by immunoblotting. As shown in Fig. 1D, with 5 out of 6 compounds the variations in DAB2IP protein levels reflected the variations in Lum/Fluo ratio, thus confirming the screening results.

**Fig. 1.**
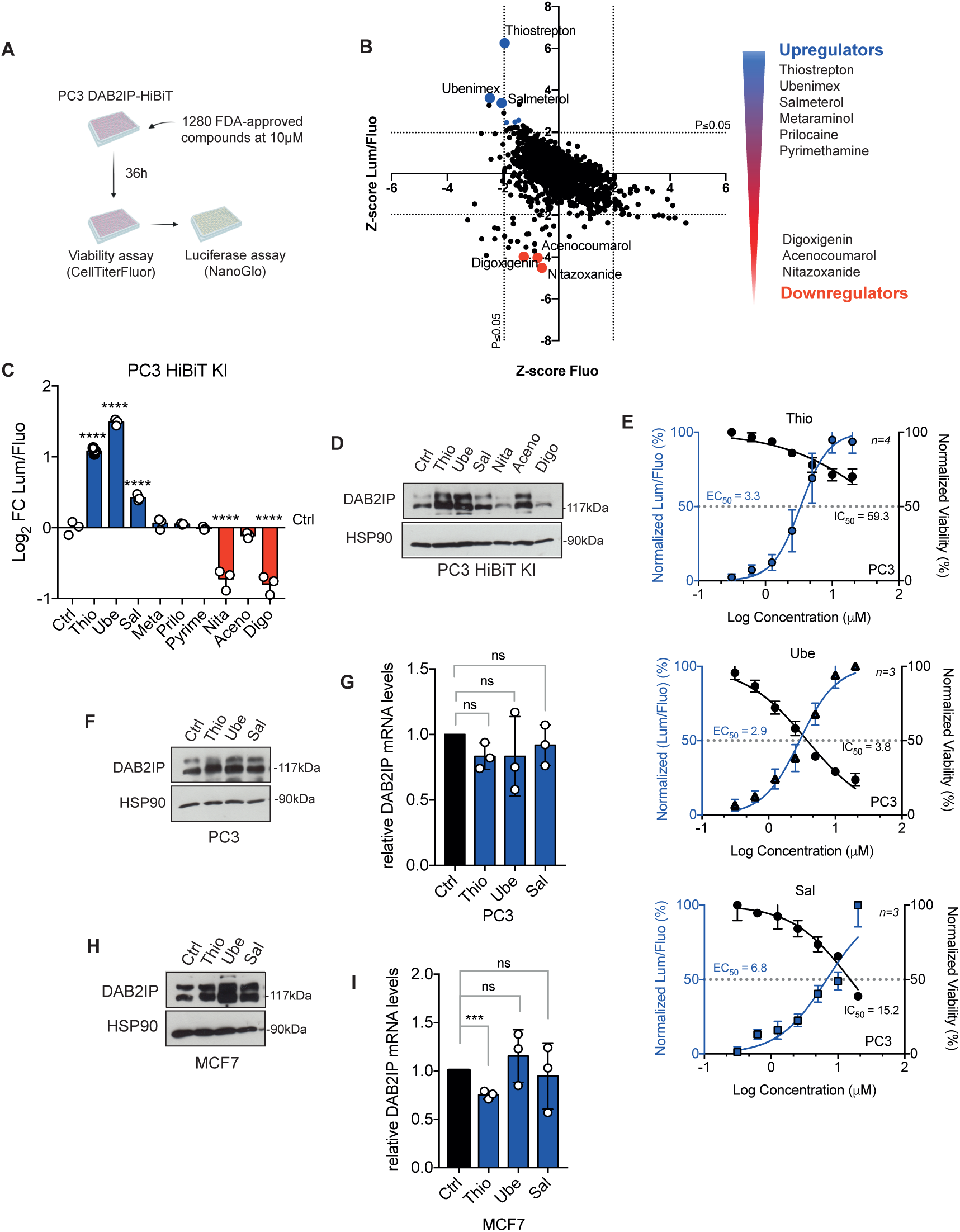
Identification of FDA-approved drugs modulating DAB2IP protein levels. **A.** Schematic representation of the high-throughput screening procedure. PC3-HiBiT cells were seeded in 384-well plates and 24 hours later 1280 FDA-approved compounds were added at 10 μM final concentration. 36 hours after treatment, cells were incubated with CellTiterFluor first, and then with NanoGlo lytic reagent. Fluorescence and luminescence readouts were acquired separately. **B.** The graph shows the Z-score of luciferase (DAB2IP levels) over fluorescence (cell viability) on the Y-axis, plotted against the Z-score of fluorescence (viability) on the X-axis. *P*-values of 0.05 were used as thresholds: hits were selected among compounds falling outside of the *p*-value on the Y-axis. On the right, list of selected drugs modulating DAB2IP levels. **C.** Histogram summarizes the Lum/Fluo ratio measured with PC3-HiBiT cells seeded in 96-well plates and treated for 36 hours with the indicated compounds at 10 μM concentration. Data are mean ± SD of three independent experiments (\*\*\*\**p* < 0.0001; one-way ANOVA with Dunnett’s post hoc). **D.** Western blot analysis of endogenous DAB2IP detected in PC3-HiBiT cells treated as in C. HSP90 was blotted as loading control. **E.** Dose-response curves of the three best candidate drugs. PC3-HiBiT cells were treated as in C, with variable drug concentrations. The graphs summarize the effect of each drug on DAB2IP levels (Lum/Fluo ratio) and the effect on viability (Fluorescence). The half-maximal effective concentration (EC50) and half-maximal inhibitory concentration (IC50) are also indicated. Data are mean of *n* independent experiments as indicated in figure ± SD. **F-G** Effects of candidate drugs on DAB2IP levels in non-edited parental PC3. **(F)** Cells were treated with the indicated drugs at 10 uM for 36h. DAB2IP was detected by immunoblotting, with HSP90 as loading control. (**G)** Cells were treated as in F. DAB2IP mRNA levels were measured by RT-qPCR. Values were normalized on histone H3 and compared to DMSO-treated controls. Data are mean ± SD of three independent experiments (ns= non-significant; one-way ANOVA with Dunnett’s post hoc). **H-I.** Effects of candidate drugs on DAB2IP in the breast cancer cell line MCF7. **(H)** Cells were treated with the indicated drugs at 10 uM for 36h. DAB2IP was detected by immunoblotting, with HSP90 as loading control. **(I)** Cells were treated as in H. DAB2IP mRNA levels were measured by RT-qPCR. Values were normalized on histone H3 and compared to DMSO-treated controls. Data are mean ± SD of three independent experiments (\*\*\**p* < 0.001, ns= non-significant; one-way ANOVA with Dunnett’s post hoc).

In the context of this study, we focused only on DAB2IP upregulators. As shown in Fig. 1E, thiostrepton (Thio) and salmeterol (Sal) displayed similar half-maximal effective concentration (EC50) values of 3.3 μM and 2.9 μM, respectively, whereas ubenimex (Ube) had an EC50 of 6.8 μM. Concerning the drug effects on viability, only ubenimex sensibly affected fluorescence, while thiostrepton and salmeterol displayed half-maximal inhibitory concentrations (IC50) significantly above the EC50 (Fig. 1E).

To exclude clone-specific artifacts, we also tested the hit compounds on parental non-edited PC3 cells, confirming the results observed in the PC3-HiBiT clone (Fig. 1F). To understand the possible mechanisms underlying drug-induced DAB2IP modulation, we investigated whether their action occurred at the transcriptional level. By RT-qPCR, we observed that all drugs did not change significantly the mRNA levels of DAB2IP, suggesting that they may act on protein synthesis or turnover (Fig. 1G).

The hit compounds were also tested on the other edited prostate cancer cell line, DU145-HiBiT (Fig. S3A-B) and on parental non-edited DU145 cells (Fig. S3C), confirming their action on DAB2IP. Of note, in DU145, Thio displayed a stronger toxicity than in PC3; nonetheless, using a lower drug concentration (4 μM) we confirmed it could increase DAB2IP protein levels also in this cell line (Fig. S3C). Similar to what observed in PC3, also in DU145 the three drugs did not significantly increase DAB2IP mRNA levels (Fig. S3D).

Finally, we tested the drugs in a different tumor type; we choose the MCF7, a luminal breast cancer cell line, estrogen receptor and progesterone receptor positive, with wild-type p53, thus representing a significantly different model with respect to PC3. Noticeably, in MCF7 the drugs displayed the same effects on DAB2IP protein levels, without significant alteration of the transcript levels (Fig. 1H-I). We conclude that Thio, Sal and Ube can increase endogenous DAB2IP protein in at least three different cell lines, representing both prostate and breast cancer.

### Thio, Sal, and Ube increase DAB2IP levels by exploiting different molecular mechanisms

The three identified drugs belong to different classes and have different established targets. Thiostrepton is thiopeptide antimicrobial drug FDA-approved for veterinary use, and known to be an inhibitor of the transcription factor FOXM1 in mammalian cells [14]. Ubenimex is a non-selective inhibitor of aminopeptidases approved in Japan for the treatment of acute non-lymphocytic leukemia under the name Bestatin, which is under clinical trials for the treatment of multiple solid tumors [15, 16]. Salmeterol is a highly selective long-acting beta-2 adrenergic agonist (LABA) that is currently prescribed for the treatment of asthma and chronic obstructive pulmonary disease (COPD) [17, 18].

To investigate whether the three drugs may have some common mechanisms of action, we used the CLUE platform (https://clue.io) to explore the Broad Institute ConnectivityMap; noticeably, very limited overlaps emerged considering both perturbagens and compounds associated to each of the drugs (Fig. S4A). We then analyzed the transcriptional effects of these drugs in PC3 and MCF7, the two cell lines in which we have functional evidence of DAB2IP upregulation, using the corresponding gene-expression data from ConnectivityMap. The three drugs have a remarkably different impact on transcription, with Thio moving a much greater number of genes than Ube and Sal; therefore, for comparative functional annotation we selected the top 1500 genes differentially affected by each drug. As shown in Fig. 2A and Fig. S4B, while there was some overlap (14-18 %) between the various drugs, only a minimal number of DEGs (2.6-2.8 %) were shared by all three, both in PC3 and in MCF7. We then used the ShinyGO platform to explore the functional annotation of genes modulated by the drugs. Gene-set enrichment analysis failed to identify obvious common pathways affected by all drugs (KEGG pathways, Fig. 2B and Fig. S4C), in line with the notion that they have different molecular targets and may act on DAB2IP via different mechanisms. Nonetheless, in both cell lines we detected enrichment for hallmarks such as TNF signaling, Hypoxia, KRAS signaling, Apoptosis, and EMT (Hallmarks.MsigDB, Fig. S5), that are related to known DAB2IP functions [2, 3].

**Fig. 2.**
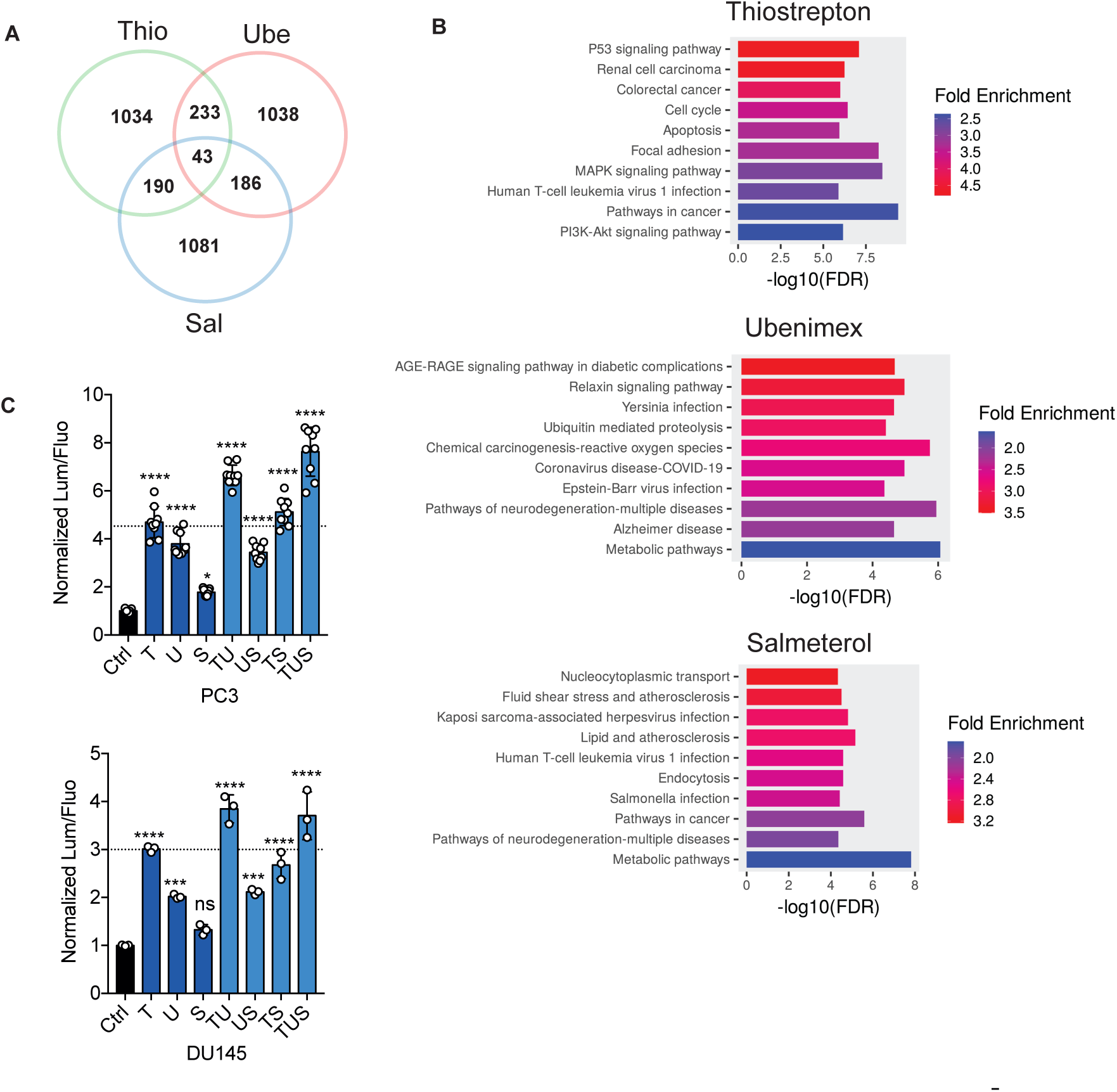
The identified drugs likely act on DAB2IP via different molecular mechanisms. **A.** Venn diagram summarizing the top 1500 genes affected by 24 hours treatment with 10 μM Thio, Ube, or Sal in PC3 cells (from ConnectivityMap). **B.** For each drug, the top 1500 genes were used for gene set enrichment analysis (KEGG pathways) using the ShinyGO platform. **C.** Thio, Ube, and Sal display additive effect on DAB2IP upregulation. PC3-HiBiT and DU145-HiBiT cells were treated for 36h with various combinations of the drugs as indicated, at a total 10μM final concentration. Endogenous DAB2IP levels were quantified as in Fig. 1C.

Finally, using the PC3-HiBiT and DU145-HiBiT cell lines, we observed greater accumulation of endogenous DAB2IP when the drugs were used in combination, a result that further supports the idea that these drugs act on DAB2IP via independent mechanisms (Fig. 2C). In the absence of an obvious common pathway triggered by the three drugs, we decided to focus on thiostrepton, since it displayed the most robust effect on DAB2IP levels.

### Thiostrepton does not increase DAB2IP levels by inhibiting FOXM1

One of the functionally most relevant targets of thiostrepton in mammalian cells is the transcription factor FOXM1, since Thio has been reported to inhibit its transcriptional activity and its expression levels [19, 20]. FOXM1 regulates the expression of a plethora of genes involved in cellular processes such as cell cycle, DNA repair, senescence, apoptosis, migration, invasion, and drug resistance [21]. We therefore asked whether FOXM1 inhibition would be sufficient to increase DAB2IP protein levels, possibly via an indirect mechanism. To this aim, we treated PC3-HiBiT cells with two unrelated FOXM1 inhibitors, FDI-6 and the natural compound Honokiol. Only Thio caused an increase in DAB2IP protein levels as measured with Lum/Fluo ratio (Fig. 3A). We also depleted endogenous FOXM1 using two different siRNAs, and again we detected no increase in DAB2IP levels (Fig. 3B). Furthermore, we confirmed these observations in non-edited PC3 and in MCF7 cells, where no increase in DAB2IP could be detected by immunoblotting upon drug-mediated FOXM1 inhibition (Fig. 3C), or siRNA-mediated FOXM1 knockdown (Fig. 3D). Together, these results strongly indicate that Thio does not affect DAB2IP levels via inhibition of FOXM1, implicating a different molecular mechanism for DAB2IP upregulation, most likely non-transcriptional, that remains to be uncovered.

**Fig. 3.**
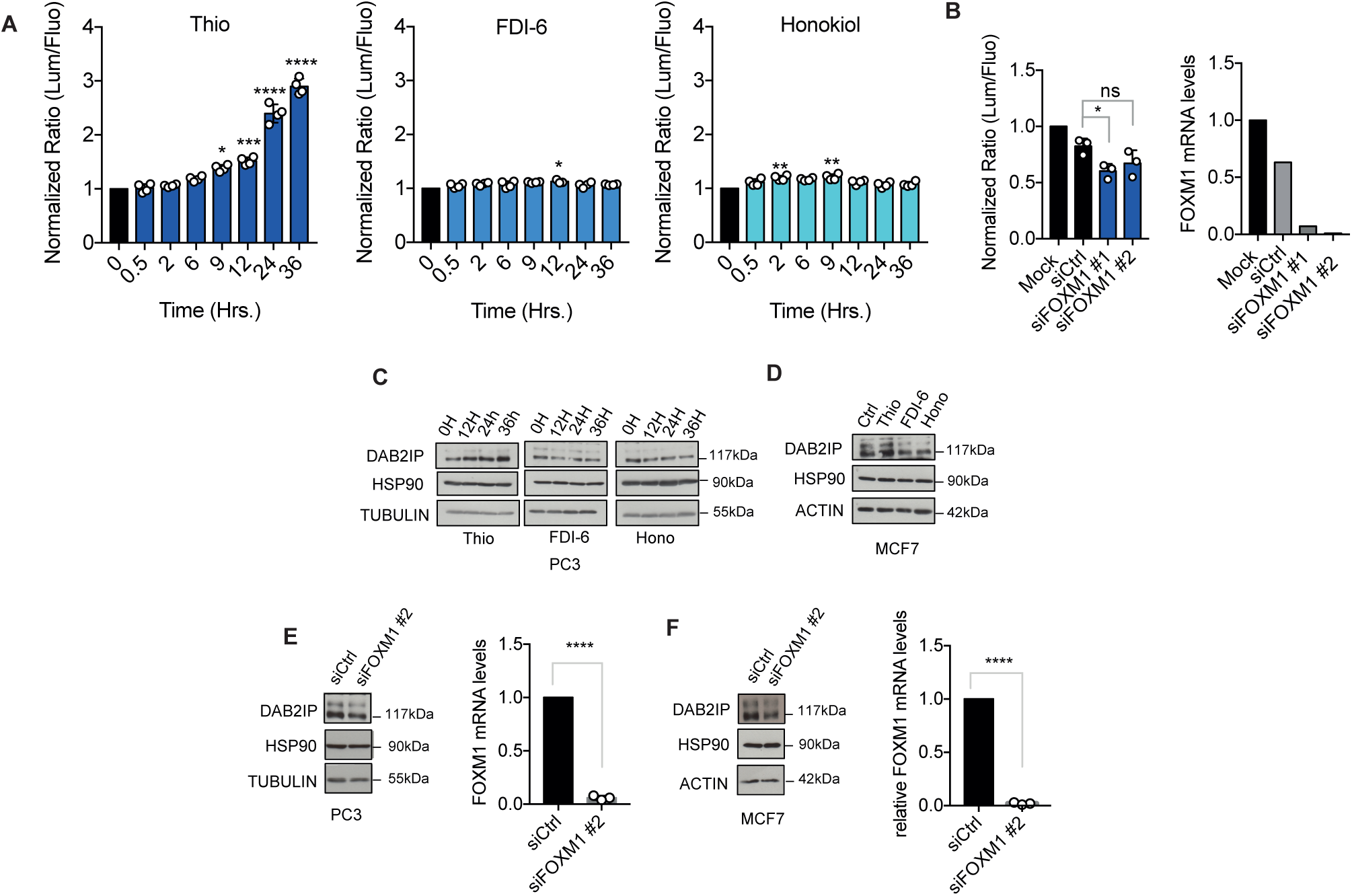
FOXM1 inhibition does not affect DAB2IP levels. **A.** Two unrelated FOXM1 inhibitors do not increase DAB2IP levels. PC3-HiBiT cells seeded in 96-well plates were treated with 10 μM thiostrepton, FDI-6, or Honokiol for the indicated time points and endogenous DAB2IP levels were measured by Luc/Fluo ratio as in Fig. 1. Data are mean ± SD of three wells per condition (\**p* < 0.1, \*\*\**p* < 0.001, \*\*\*\**p* < 0.0001; one-way ANOVA with Dunnett’s post hoc). **B.** FOXM1 silencing does not increase DAB2IP levels. Left, PC3-HiBiT cells were transfected for 48 hours with two different siRNAs targeting FOXM1, and DAB2IP levels were measured by Luc/Fluo ratio as in A. Right, FOXM1 mRNA levels were measured by RT-qPCR as a control of silencing efficiency. Data are mean ± SD of three wells per condition (one-way ANOVA with Dunnett’s post hoc). **C.** Parental non-edited PC3 cells were treated as in A. Endogenous DAB2IP protein was detected by immunoblotting with HSP90 and tubulin as loading controls. **D.** MCF7 cells were treated for 36 hours with the indicated inhibitors at 10 μM concentration. Endogenous DAB2IP protein was detected by immunoblotting, with HSP90 and actin as loading controls. **E.** Parental non-edited PC3 cells were transfected for 48 hours with FOXM1 siRNA. Left, representative western blot of endogenous DAB2IP. HSP90 and tubulin were blotted as loading controls. Right, FOXM1 mRNA levels measured by RT-qPCR as a control of silencing. **F** MCF7 cells were treated and analyzed as in E. Left, representative western blot of endogenous DAB2IP. HSP90 and actin were blotted as loading controls. Right, FOXM1 mRNA levels measured by RT-qPCR as a control of silencing. Data are mean ± SD of three independent experiments (\*\*\*\**p* < 0.0001; unpaired Student’s t-test).

### The cancer-inhibitory effects of thiostrepton are mediated in part by DAB2IP upregulation

To investigate the impact of thiostrepton on oncogenic phenotypes, we performed proliferation, migration and invasion assays with PC3 cells. To minimize nonspecific effects due to toxicity, Thio was tested at its EC50. Despite the relatively low concentration, it effectively reduced the proliferative capabilities of prostate cancer cells, as assessed by colony formation assay (Fig. 4A). Moreover, it significantly decreased the migratory and invasive capabilities of PC3, as confirmed by wound healing and matrigel invasion assays, respectively (Fig. 4B-C).

**Fig. 4.**
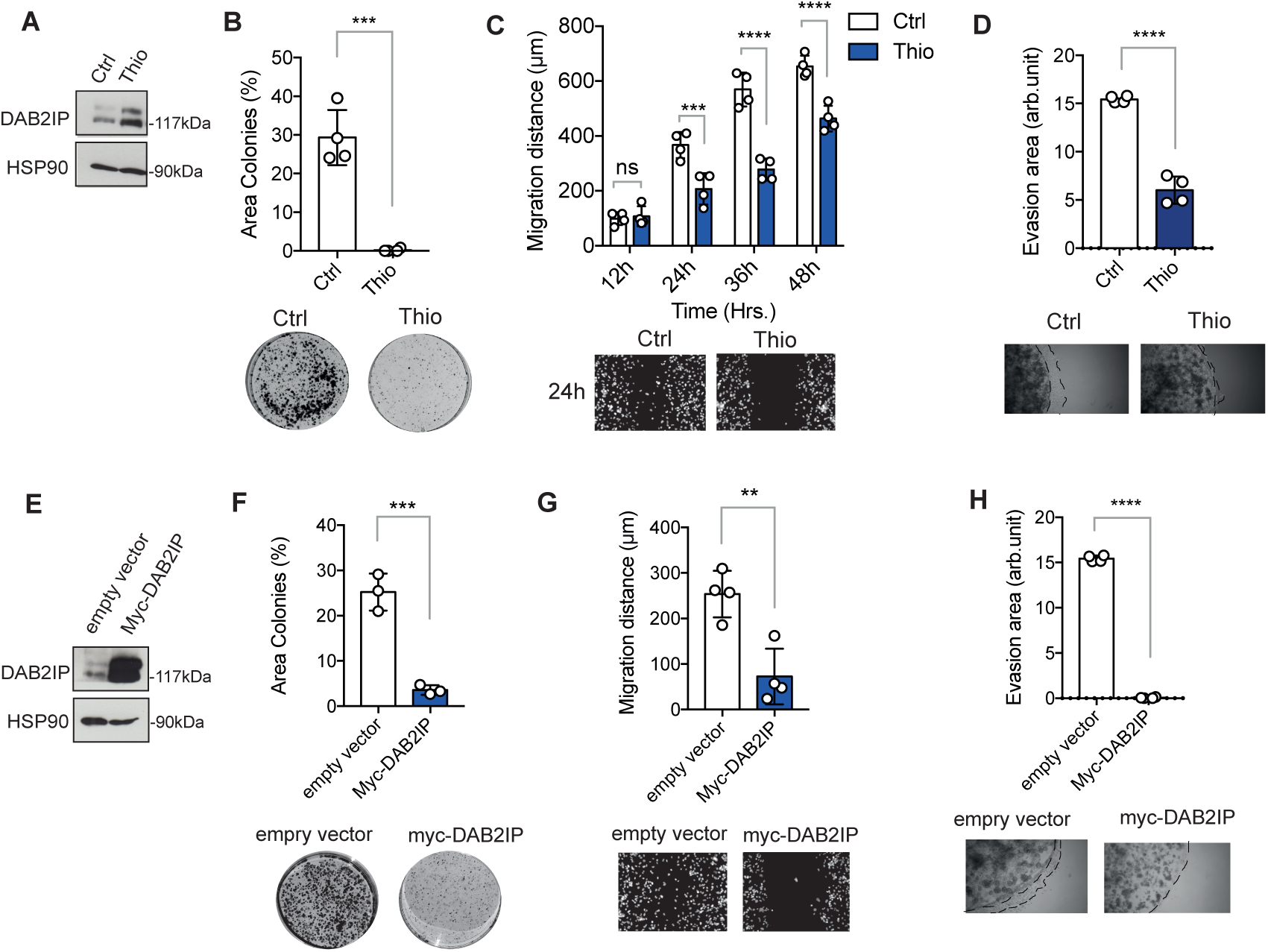
Treatment with thiostrepton inhibits aggressive phenotypes of prostate cancer cells in vitro, an effect similar to DAB2IP overexpression. **A.** Representative immunoblotting of PC3 cells untreated or treated with 3 μM Thio for 48 hours, with HSP90 blotted as loading control. **B.** Thio inhibits colony formation. An identical number of PC3 cells were plated at low density and treated with 3 μM Thio or DMSO (Ctrl) for 48h; after 10 days, colonies were stained and photographed (representative pictures are shown). Colony formation efficiency was quantified using ImageJ (cells covered area/dish area). Results are mean ± SD of four indipendent experiments (unpaired Student’s t-test). **C.** Thio reduces cell migration. PC3 cells were grown until 90% confluence, then a central region was scraped off with a sterile tip and cells were treated with 3 μM Thio or DMSO (Ctrl). Wound closure was checked at the indicated time-points (representative images at 24 hours are shown). Migration distance was quantified using ImageJ as indicated in materials and methods. Results are mean ± SD of four wells per condition (two-way ANOVA with Sidak correction). **D.** Thio inhibits cell evasion in matrigel. Cells were pre-treated for 24 hours with 3 μM Thio or DMSO (Ctrl). Then, the same number of cells were seeded inside a matrigel drop in 0.1% serum covered with 10% serum medium. Evasion events were monitored during the following days by phase contrast microscopy (representative images at 7 days are shown). Graph summarizes the area of evasion events quantified using ImageJ and expressed as arbitrary units. Results are mean ± SD of four indipendent experiments (unpaired Student’s t-test). **E.** Representative immunoblotting of control and DAB2IP-stably expressing PC3 cells, with HSP90 blotted as loading control. **F.** DAB2IP over-expression inhibits colony formation. An identical number of PC3 cells stably expressing myc-DAB2IP or empty vector were plated at low density; colony formation efficiency was quantified as in A. **G.** DAB2IP over-expression reduces cell migration. PC3 cells stably expressing myc-DAB2IP or empty vector were plated at high confluence and wound-healing assays were performed and analyzed exactly as in B. **H.** DAB2IP inhibits cell evasion in matrigel. PC3 cells stably expressing myc-DAB2IP or empty vector were seeded in Matrigel drops, and cell evasion events were monitored and quantified as in C. (\*\**p* < 0.01, \*\*\**p* < 0.001, \*\*\*\**p* < 0.0001; unpaired Student’s t-test).

To compare the effects of thiostrepton with those of DAB2IP up-regulation, we stably expressed DAB2IP in PC3 cells. In line with several previous observations, ectopic DAB2IP strongly inhibited colony formation, cell migration, and matrigel invasion (Fig. 4D-F).

To investigate the role of DAB2IP upregulation in the onco-suppressive effects of thiostrepton, we depleted DAB2IP in PC3 using two different sgRNAs, and analyzed the effects of the drug in these cells. Strikingly, DAB2IP loss dampened the inhibitory effect of Thio on cell proliferation, as assayed by colony formation (Fig. 5B). Similarly, DAB2IP loss mitigated the inhibitory effect of Thio on cell migration in wound-healing assays (Fig. 5C).

**Fig. 5.**
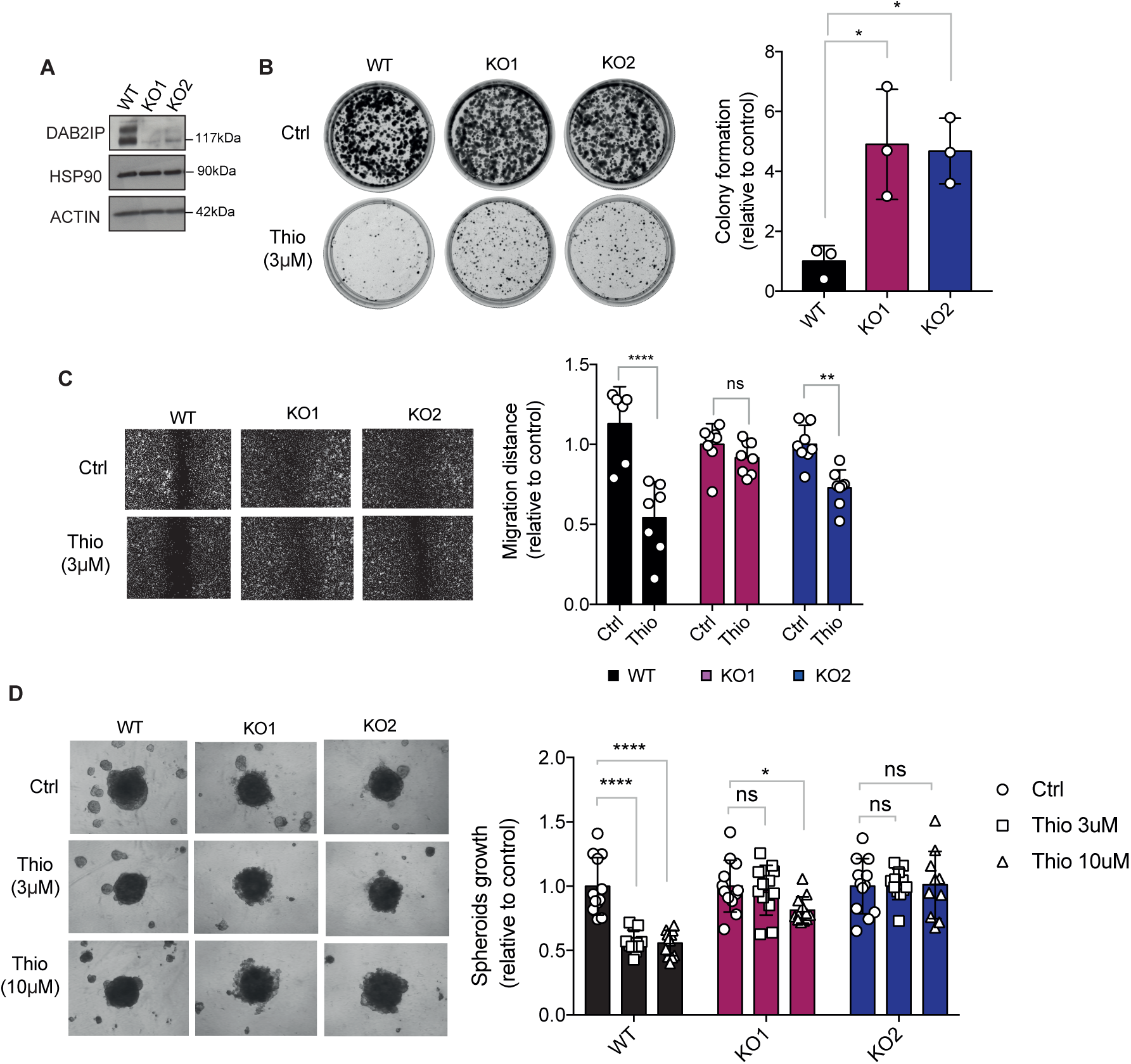
The inhibitory effects of thiostrepton on cancer cells are reduced upon DAB2IP depletion. **A.** Western blot analysis of DAB2IP expression in parental (WT) and DAB2IP knock-out (KO1, KO2) PC3 cells, with HSP90 and actin as loading controls. **B.** Thiostrepton is less effective in suppressing colony formation in the absence of DAB2IP. An identical number of PC3 cells (WT, KO1, KO2) were plated at low density. After 24 hours cells were treated with 3 μM Thio or DMSO (Ctrl) for 48h, and then cultured for 14 days. Colonies were stained, photographed (representative pictures are shown), and colony formation efficiency was quantified as in Fig. 4A. The graph summarizes colony formation in treated samples normalized to their respective controls (DMSO). Results are mean ± SD of three indipendent experiments (\**p* < 0.1; one-way ANOVA with Dunnett’s post hoc). **C.** Thiostrepton is less effective in suppressing cell migration in the absence of DAB2IP. PC3 cells (WT, KO1, KO2) were grown until 90% confluence, then a central region was scraped off with a sterile tip and cells were treated with 3 μM Thio or DMSO (Ctrl). Wound closure was checked at 48 hours (representative images are shown). Migration distance was quantified using ImageJ as indicated in materials and methods, and normalized to the respective DMSO controls. Results are mean ± SD of eight wells per condition (\**p* < 0.1, \*\*\**p* < 0.001, ns = non-significant; two-way ANOVA with Sidak correction). **D.** Thiostrepton is less effective in suppressing growth of prostate cancer cell spheroids in the absence of DAB2IP. An identical number of cells (WT, KO1, KO2) were plated in 96-well ultra-low attachment plates to form spheroids. After 48h, spheroids were treated with thiostrepton at the indicated concentrations or DMSO (Ctrl). The area of spheroids was measured 7 days after treatment (representative images are shown). Results are mean ± SD of three indipendent experiments (n=4 or 6 spheroids per replicate; \**p* < 0.1, \*\**p* < 0.01 \*\*\*\**p* < 0.0001, ns = non-significant; two-way ANOVA with Tukey correction).

Finally, we investigated the effects of Thio on the capability of PC3 to grow into spheroids. Consistently with previous assays, Thio strongly reduced spheroid growth of parental PC3. Despite the fact that knock-out cells form spheroids that are morphologically distinct from WT cells, it is evident that Thio had a much milder effect on 3D growth of DAB2IP KO cells (Fig. 5D). Taken together, these data indicate that upregulation of DAB2IP protein contributes to the cancer inhibitory action of Thio, and may have potential implications for the clinical repurposing of this drug.

## Discussion

The tumor suppressor DAB2IP is frequently downregulated in various human malignancies by multiple mechanisms, but is rarely deleted or mutated; this makes it a good candidate for development of anti-cancer drugs that increase its expression. In this drug-repurposing screen, we searched for molecules able to modulate, both positively and negatively, the expression levels of endogenous DAB2IP. Drugs that induce DAB2IP downregulation are not interesting for their potential clinical application, but could reveal novel mechanisms of DAB2IP downregulation, that in turn may be targeted to promote DAB2IP stabilization in cancer.

According to our experience, endogenous DAB2IP cannot be reliably detected by immunofluorescence or by other imaging techniques amenable to high-throughput. Therefore, we tagged endogenous DAB2IP with the HiBiT peptide for luciferase detection. The choice of the C-terminal insertion site was driven by the need to tag all possible isoforms with minimal impact on the structure and function of the protein. With such design, the HiBiT-derived luminescence theoretically matches the total amount of expression of all possible DAB2IP isoforms.

Such an approach allows the identification of compounds that affect endogenous protein levels under native conditions, potentially acting on all the steps of its regulation, from transcription, to translation, to RNA stability and protein turnover, with high sensitivity.

Primary and secondary screening consistently identified 3 molecules that upregulate endogenous DAB2IP in various cell lines. None of these compounds altered DAB2IP mRNA levels, suggesting a post-transcriptional or post-translational mechanism of action. In this regard, DAB2IP has been reported to be downregulated by E3 ubiquitin-ligases Skp2 and Smurf1 [22, 23] and is subject to translational inhibition by several microRNAs [7, 24–26]. It is therefore possible that similar mechanisms are targeted by some of the identified compounds, however additional studies are needed to further dig into the molecular mechanisms underlying the action of these drugs.

Of the 3 drugs, only ubenimex had been previously reported to increase DAB2IP levels in a mass spectrometry-based, high-throughput study of drug effects in colon cancer cells (DeepCoverMOA, [27]).

For its robust effect on DAB2IP protein levels, we focused on thiostrepton, a thiopeptide antibiotic that has been reported to have anticancer activity in vitro, supporting its possible repositioning for cancer treatment. In this regard, there are abundant preclinical evidence describing the effect of thiostrepton in inhibiting proliferation and survival of various tumor cell lines [28–31]. Despite this, it is not yet approved by FDA for cancer treatment.

Our experiments confirmed that thiostrepton counteracts aggressiveness of prostate cancer cells in vitro, but also revealed that this effect is less efficient in the absence of DAB2IP, thus suggesting that DAB2IP upregulation contributes to the anti-tumoral action of the drug.

The anti-cancer properties of thiostrepton have been essentially ascribed to its inhibition of the Forkhead box M1 (FOXM1) transcription factor, that regulates cell cycle genes essential for DNA replication and mitosis [20, 32, 33].

We have collected evidence to reasonably exclude that DAB2IP upregulation is the indirect consequence of Thio-mediated FOXM1 inhibition, so the two proteins can be considered as independent targets of the drug. In this regard, it is legitimate to speculate that increased DAB2IP levels can support the anti-tumoral effects of FOXM1 inhibition in certain cell contexts, or can mediate cytostatic effects of Thio in cancer cells that are not dependent on FOXM1 activity.

In conclusion, this study supports the concept that DAB2IP can be a druggable molecule, and its drug-induced upregulation, by thiostrepton treatment, can counteract different pro-oncogenic features in prostate cancer cells. Together, these findings set the basis to explore a possible repositioning of thiostrepton for the treatment of prostate cancer, unveiling DAB2IP as an important factor for its tumor-suppressive activity.

Finally, from a methodological perspective, our work demonstrates the applicability of the HiBiT system for high-throughput quantitation of an endogenous protein, indicating it as an efficient tool for drug discovery. This kind of approach might be applied to screen, in a reliable and convenient way, molecules that increase or decrease the endogenous levels of any protein of interest, in particular those that are not easily detected by other techniques.

## Materials and Methods

### Cell Lines, Transfections and Drug Treatments

PC3 were cultured in DMEM-F12 (1:1) (ECM0728L, ECM0135L, Euroclone) supplemented with 10% FBS, pen/strep solution, MEM Nonessential Aminoacids (ECB3054D, Euroclone), and Sodium Pyruvate (1 mM), (ECM9542D, Euroclone). DU145 were cultured in EMEM (LOBE12611F, Euroclone), supplemented with 10% FBS and pen/strep solution. MCF7 were cultured in EMEM (LOBE12611F, Euroclone), supplemented with 10% FBS, pen/strep solution, and human insulin 10ug/ml (Sigma, I2643). H1299 cells were cultured in RPMI (ECM2001L, Euroclone) supplemented with 10% FBS, and pen/strep solution. PC3, DU145 and MCF7 cells were purchased from ATCC. Cells were cultured at 37°C in a humidified incubator with 5% CO_2_. All cell lines were negative for mycoplasma contamination.

For knockdown experiments, cells were transfected the same day of plating (384-well plates) or 24 hours after plating (6-well plates) with 50nM siRNA (384-well plates) or 40nM siRNA (6-well plates) or 3nM miRNAs. Cells were processed after 48 hours, except when differently specified. DAB2IP silencing was performed using a mix of siDAB2IP A and siDAB2IP B.

Cells were treated with FDA-approved drugs included in the Prestwick Chemical library, that were purchased from GreenPharma. For validation experiments, cells were treated with thiostrepton CAS-1393-48-2-Calbiochem (Sigma-Aldrich). In all experiments cells were treated with drugs or with an equivalent volume of DMSO for 36 hours, unless otherwise indicated.

### High-Throughput Screening

The screening was performed using a clone of PC3-HiBiT cells with homozygous insertion of the tag. PC3-HiBiT cells (4.0×10^3^ per well) were seeded in white opaque 384-well microplates [Perkin Elmer, 6007690], 24 hours later 1280 FDA-approved & EMA-approved drugs [Prestwick Chemical Library®] were transferred robotically from library stock plates (1mM in DMSO) to the plates containing the cells; control (DMSO) was added to columns 1, 2, 23 and 24 of each plate. Cells were processed 36h after addition of drugs. Briefly, wells were aspirated with Plate washer BioTek 405, leaving 10 μl of medium. 10 μl of 2X CellTiterFluor reagent (Promega) was added to the cells, and cells were incubated for 30 min at 37°C, 5% CO2. Fluorescence was detected using EnVision multimode plate reader with dual monochromator [PerkinElmer] (400nmEx/505nmEm). Immediately after, 20 μl of 2X NanoGlo Lytic reagent (Promega) was added to the wells. Cells were incubated for 10 min at room temperature and luminescence was detected with EnVision multimode plate reader. Luminescence values were normalized on fluorescence readings. Screening was performed once, at 10 μM drug concentrations; final concentration of DMSO in the culture medium was 1% (v/v).

### Gene knock-out

For DAB2IP knock-out, sgRNAs were designed following previously described protocol [13]. PC3 cells were nucleofected using a NEPA21 electroporator (NepaGene). Cells were washes twice with PBS to remove serum components and were resuspended in Opti-MEM (Life Technologies, 31985047). 6.0×105 cells per reaction were dispensed in electroporation cuvettes (NEPA EC-002S, 2 mm gap) in a total final reaction volume of 70 μl. For each reaction 2,4 μl of Cas9 Electroporation Enhancer (IDT) and 10 μl of RNP complex were added. RNP complex was prepared as previously described [13]. Opti-MEM was added to the mixture to reach the final reaction volume. Poring pulse was 175 V for 2.5 msec pulse length twice with a 50 msec interval between the pulses and 10% decay rate with + polarity. The transfer pulse condition was five pulses at 20 V for 50 msec pulse length with 50 msec interval between the pulses and 40% decay rate with +/− polarity. Immediately after electroporation cells were recoverd from the cuvette and dispensed into a medium filled 6-wells plate. Efficiency of gene editing was evaluated by analyzing Sanger sequencing results of PCR products of target genomic region using TIDE software. Primers used for genomic amplification are listed in Table 3.

**Table 1.**
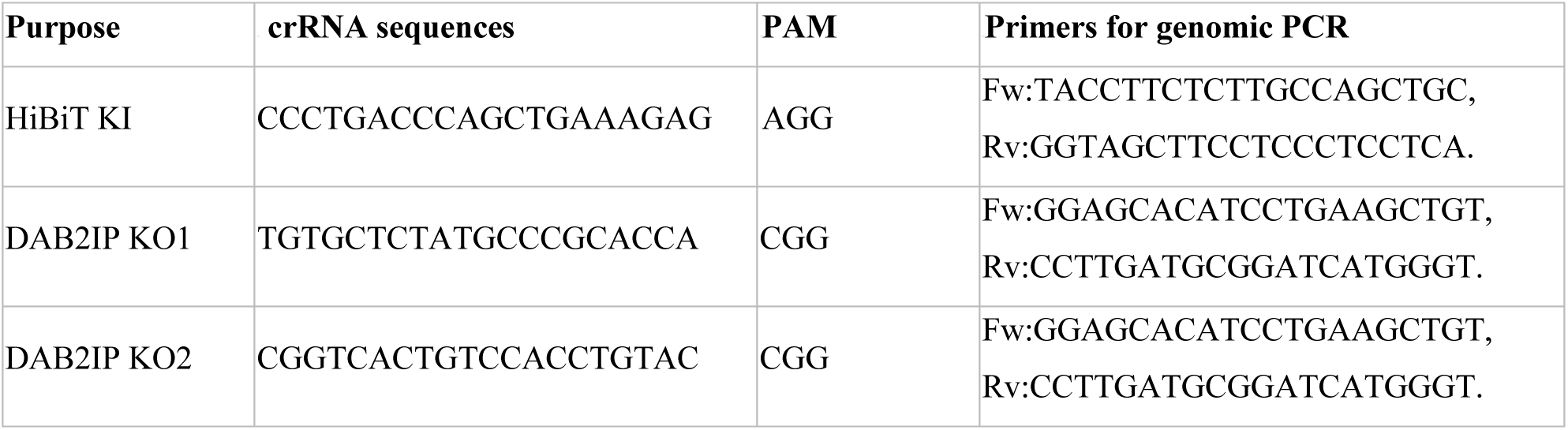
Sequences of crRNAs used:

**Table 2.**
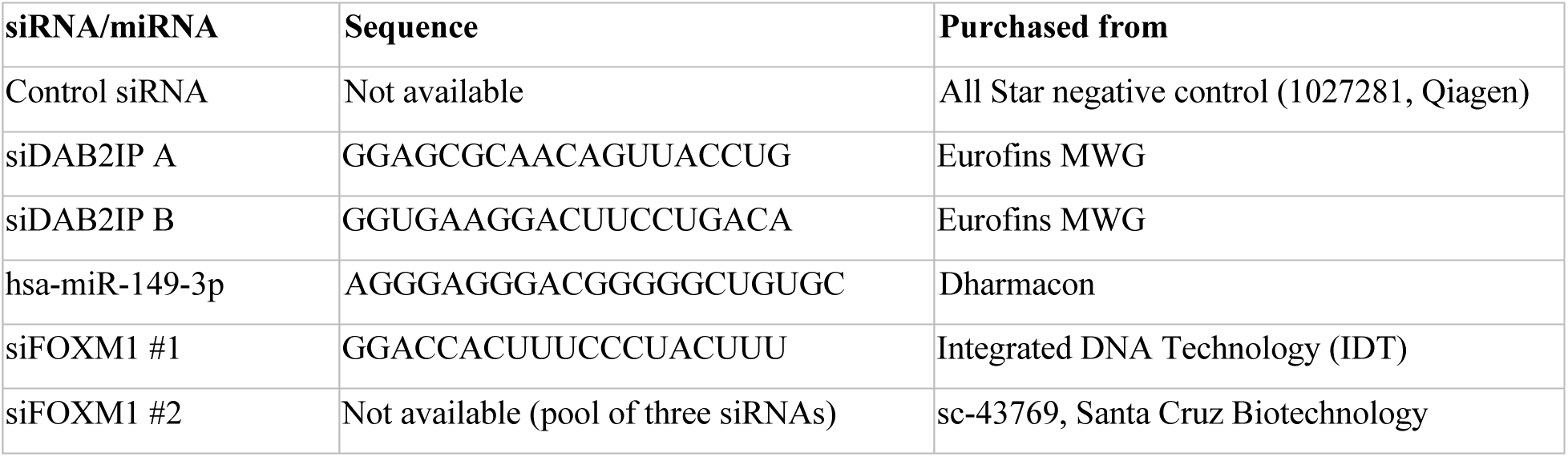
Sequences of siRNAs/miRNA used:

**Table 3.**
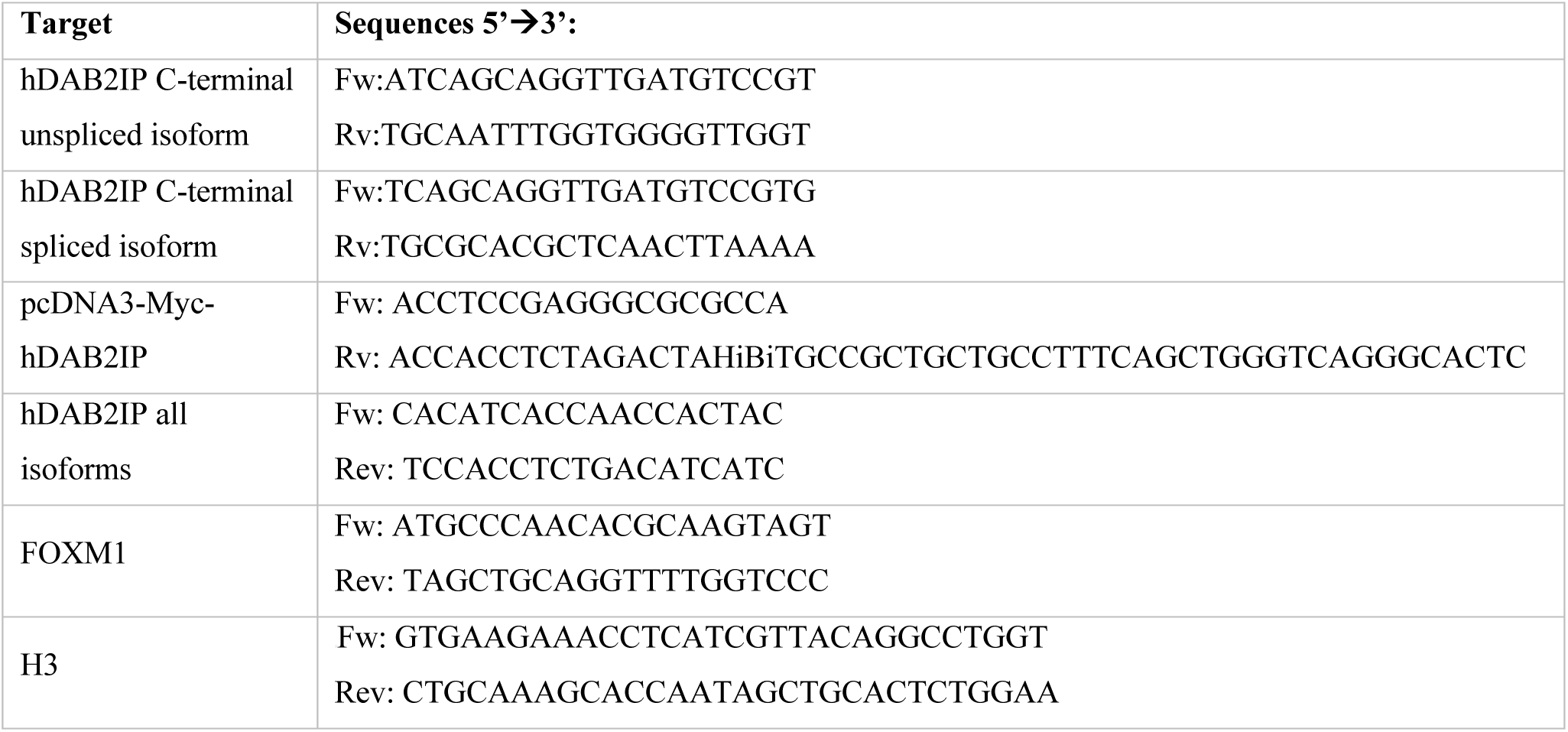
Sequences of primers used:

### Protein Expression Analysis

Cells were lysed in 2X SDS Sample buffer (125 mM Tris-HCl at pH 6.9, 4% SDS, 20% Glycerol, 3% β-mercaptoetanol). Samples were sonicated and boiled at 95°C for 5 min. About 10-20 ug of lysates were loaded in polyacrylamide gels and resolved by SDS-PAGE (BioRad Mini-Protean® Tetra System) and transferred to nitrocellulose membranes [GE Healthcare, 10600001]. Membranes were blocked for at least 1 hour at room temperature in 5% skim milk PBST (0,1% Tween-20 in PBS) and incubated overnight at 4°C with primary antibodies diluted in 5% skim milk PBST (Table 4). Membranes were washed three times in PBST for 10 min. Then were incubated for 1 hour at 4°C with horseradish peroxidase (HRP)-conjugated secondary antibody diluted 1:4000 in 5% skim milk PBST. Membranes were washed as above and rinsed in PBS before detection. Protein detection was performed by chemiluminescence using Liteablot Extend [EMP013001, Euroclone] for DAB2IP and c-Myc, and with ECL [32209; Thermo Scientific] for normalization markers. Chemiluminescent signal was impressed on Hyperfilm ECL [Amersham].

**Table 4.** List of primary antibodies used:

| Antigen | Species | Company | Dilution |
| --- | --- | --- | --- |
| DAB2IP | Rabbit (polyclonal) | Bethyl, A302-440A | 1:4000 |
| Myc-tag | Mouse (monoclonal) | 9E10 hybridoma supernatant | 1:100 |
| HSP90 | Mouse (monoclonal) | Santa Cruz, sc-13119 | 1:8000 |
| Actin | Rabbit (polyclonal) | Sigma, A2066 | 1:8000 |
| Tubulin | Mouse (monoclonal) | Sigma, T5168 | 1:8000 |

### RNA Expression Analysis

Total RNA was extracted with TRIFAST II [EMR517100, Euroclone] using a standard protocol. For RT-qPCR, 500 ng of total RNA was reverse transcribed with iScript Advanced cDNA Synthesis Kit (mRNA) [1725038, Biorad]. Real-time PCR was performed using iTaq Universal SYBR Green SMX 5000 [1725124, Biorad] on a CFX96 Real-Time PCR System [Biorad].

### Colony Formation Assay

Cells were seeded at a density of 5000 cells per 6-cm diameter plate and incubated for 48 hours in complete culture medium, then cells were treated with drugs at EC50 concentration for 48 hours. After 10-14 days, cells were fixed and stained with 10% MetOH and 0.1% crystal violet [C-6158, Sigma] in H20 for 30 min. Plates were photographed and colony formation efficiency was calculated by measuring the % of colonies-covered area over the whole dish area. The areas were measured using ImageJ software.

### Wound Healing Assay

Cells were plated on 96-well plates in presence of Hoechst (0.2 μg/ml) [LifeTech] and cultured to 90% confluence. Cells were scraped with a sterile pipette tip, and treated with drugs (or DMSO) at EC50 concentration. The scratch area was photographed at defined time points, and cell migration was quantified and expressed as average rate of closure of the scratch. Images of live cells were automatically acquired with PerkinElmer Operetta every 12 hours. The width of the scratches was measured using ImageJ. Migration distances, M(t), were calculated as followed: M(t) = width (0) – width (t), where width (t) is the wound width at time t and width (0) is its initial width.

### Matrigel Drop Invasion Assay

PC3 cells were pre-treated with indicated drugs for 48h. Then 7500 cells per condition were plated in 12 μl of BD Matrigel (7mg/ml) [BD Bioscience] in low serum (0.1% FBS). Five drops of 12 μl of Matrigel with cells were plated for each condition and then coated with 2 ml of high serum (10% FBS) medium. After 7 days, the drops were fixed and stained with 10% methanol and 0.1% crystal violet in H20. Evasion was measured using ImageJ software on 5 random non-overlapping microscope fields at 100X magnification of four different drops for each condition.

### Tumor Spheroid Assay

500 cells/well were seeded in ultra-low attachment 96-well plates round bottom in complete medium plus 2,5% matrigel as previously described [34]. After 48 hours spheroids were treated with 3 μM or 10 μM of thiostrepton or DMSO as control. Images of live spheroids were taken at the time of treatment (t0) and after 7 days (t) using ZEN Microscopy Software (ZEISS) at 100X magnification. Area of spheroids was measured in arbitrary units using ImageJ software on 4-6 spheroids per condition. Spheroids growth was calculated as follows: Area (t)/Area (t0).

### Analysis of ConnectivityMap datasets

Gene-expression data for PC3 and MCF7 treated with the various drugs, reported as weighted Z-scores, were downloaded from CLUE (https://clue.io). We then used the expression vector, calculated by computing the Spearman correlation among replicates as reported in the CLUE guidelines, to identify differentially expressed genes (DEGs). We considered significant all the genes with adjusted *p*-value lower than 0.05. To compare the transcriptional impact of the different drugs, we sorted DEGs based on the absolute value of log fold change, and analyzed the top 1500. For gene-set enrichment analysis, we used the ShinyGO platform (version 0.80, http://bioinformatics.sdstate.edu/go/).

### Statistical Analysis

All the results are expressed as mean ± SD. Values of *p* < 0.05 were considered statistically significant. For statistical comparison of two groups, unpaired two-tailed Student’s t-test was used; for the comparison of three or more groups, one-way ANOVA followed by Dunnett’s post-hoc test, or two-way ANOVA, were used as indicated in figure legends. The number (*n*) of independent experiments is indicated in the figures or figure legends. Data were analyzed using Prism 7.0 (GraphPad).

## Acknowledgements

The authors acknowledge the contribution of Lorenzo Balbi who helped with some of the experiments, and all other members of the LC and LB groups for discussion and support. The screenings and several other quantitative analyses were performed at the ICGEB High-Throughput Screening Facility (https://www.icgeb.org/high-throughput-screening-equipment/).

## Author contributions

Conceptualization, RDFF, LLF, LB, LC. Investigation, RDFF, SM, RK, VP, SG, DS. Specific resources, LLF, LB. Supervision, LLF, LB, LC. Writing—original draft, RDFF, LC; writing— review and editing, RDFF, LLF, LB, LC. All authors have read and agreed to the current version of the manuscript.

## Ethics

Not applicable

## Data Availability

Not applicable

