## Supplemental methods and figures for "Drug-repurposing screen identifies thiostrepton as a novel regulator of the tumor suppressor DAB2IP"

### Supplementary Materials and Methods

### Supplementary figure legends

### Supplementary figures

### Supplementary Materials and Methods

#### Analysis of hDAB2IP isoforms

For assessing the expression of predicted C-terminal splice isoforms of hDAB2IP, 500 ng of total RNA was reverse transcribed with iScript Advanced cDNA Synthesis Kit (mRNA) [1725038, Biorad]. PCR reactions were done using Taq DNA Polymerase [201203, Qiagen] on a MiniAmpTM Thermal Cycler [Thermo Fisher]. PCR products were analyzed by agarose gel electrohoresis. Primer sequences are listed in Table 3.

#### DAB2IP-HiBiT plasmid generation

To evaluate the applicability of the HiBiT tag (Promega) in detecting DAB2IP protein levels we cloned HiBiT in fusion to hDAB2IP in an expression vector. To generate a plasmid encoding the DAB2IP-HiBiT protein, a C-terminal fragment of DAB2IP in fusion with the HiBiT tag was generated by PCR and inserted in the pcDNA3-Myc-hDAB2IP plasmid (kindly provided by Sidney Yu, University of Hong Kong). Transfection of pcDNA3-hDAB2IP-HiBiT in H1299 was performed using Lipofectamine 2000 (Invitrogen), following manufacturer’s instructions.

#### Retroviral Transduction

For generation of prostatic cancer cells stably expressing hDAB2IP, low confluence (~20%) 293GP packaging cells, stably expressing retroviral gag and pol proteins, were transfected by calcium-phosphate precipitation. Briefly, cells were plated the day before the transfection in 10 cm-diameter dishes in DMEM with 10% FBS and antibiotics. Each different virus was produced by co-transfection of 10 μg of the vector of interest and 5 μg of pEnv encoding vector. After 8 hours, medium was changed and cells were incubated at 37°C. After 48 hours the virus-containing medium was filtered (0.45 μm filter) and supplemented with 10% FBS and polybrene (8 μg/ml). Target cells growing at low confluence (~ 30-40%) were treated with the appropriate viral supernatant for 24 hours, and subsequently selected with puromycin (0.5 μg/ml). Single clones with the lowest levels of ectopic DAB2IP protein were selected from the bulk of infected cells and one such clone was used to perform phonotypic assays. pLPC-Myc-DAB2IP was obtained by cloning into the pLPC vector the insert from pcDNA3-Myc-hDAB2IP.

**Endogenous DAB2IP Tagging**

All synthetic nucleic acids were purchased from Integrative DNA Technologies (IDT). For CRISPR-Cas9 knock-in gene tagging, crRNAs were designed using multiple online tools, providing as search base an 80–100 bp genomic sequence centered around the preferred editing site.

To perform HDR-driven gene editing, a 150 bp ca. single stranded donor DNA (Ultramer® DNA Oligos, IDT) was synthesized containing two flanking regions (50 bp each) providing left and right homology arms, and a central sequence for the desired tag. The knock-in sequence encoded a short flexible linker (Gly-Ser) followed by the HiBiT sequence. Single stranded DNA donor was designed using an online tool (https://horizondiscovery.com/en/dharmacon) with the insertion site as close as possible to the predicted editing site. Cas9 was delivered as recombinant protein (120 pmol) together with crRNA (150 pmol) and tracrRNA (150 pmol) [Alt-R® CRISPR-Cas9 tracrRNA, IDT, 1072533] and single stranded donor DNAs (120 pmol) as previously described {Ghetti, 2021 #17}. The mixture was prepared in 16-well Nucleocuvette StripsTM [Lonza]. Cells were electroporated using the 4D-NucleofectorTM [Lonza, AAF-1002B] using the following programs: DS-137 for PC3, CA-137 for DU145. 10 min after electroporation, 75 µl of complete medium with NU7441 (1µM) were added to nucleofected wells and cells were transferred into new 12-well plate to allow for recovery. 48 hours post- nucleofection the culture medium was removed and replaced with complete medium without drug.

To obtain genetically homogenous edited clones, nucleofected cells were seeded with a density of 0.5-1 cell/well in a 96-well plate previously filled with 200 µl/well of complete medium. Cells were incubated for at least 2 weeks before characterization. At least 10 isolated monoclonal lines per condition were characterized by immunoblotting and genomic PCR.

In order to assess the allelic status of HiBiT positive clones, the integration was verified by Sanger sequencing of PCR products of target genomic region. Primers used for genomic amplification are listed in Table 3.

#### Supplementary figure legends

**Fig. S1 Characterization of human DAB2IP isoforms and tagging site.**

**A-B** Human DAB2IP is expressed with two C-terminal variants in multiple cell lines. **(A)** Schematic representation of predicted C-terminal ends with primers used to detect the unspliced (above) (NM_138709.2) or the spliced (below) isoforms (NM_032552.3). Lines represent introns, and boxes exons. Filled boxes represent the translated coding sequence. **(B)** Multiple normal and transformed cell lines express both C-terminal variants. Total RNA extracted from the indicated cell lines was reverse-transcribed and amplified using primers designed to distinguish alternatively spliced DAB2IP isoforms. In THP-1 monocytic leukemia cells none of the isoforms could be detected, in line with the notion that DAB2IP is not expressed in blood . Note: some electrophoresis images are cropped. **C.** Schematic representation of DAB2IP isoforms, with a blow-up of the tag insertion site. Greyed area, the C-terminal genomic sequence of human DAB2IP; exon sequence in uppercase, intron in lowercase. Underlined is the guide recognition site in the target sequence, with the PAM sequence in yellow. The vertical dotted line indicates the predicted cut site. The insertion site is arbitrarily positioned in correspondence of the cut site at junction between codons. In the lower part, codons (above) and the corresponding translation (below) of the insert are reported. Red capital letters refer to a flexible Gly/Ser linker followed by the HiBiT-tag (in blue) and two stop codons (in black). The donor DNA also included two homology arms (HA) of 50 bp (adapted from {Ghetti, 2021 #17}. **D.** H1299 cells were transfected with pcDNA3 myc-hDAB2IP-HiBiT plasmid, encoding a fusion protein identical to the predicted endogenous tagged product. An empty pcDNA3 plasmid was transfected as a negative control. 24 hours after transfection, variable numbers of transfected cells were processed for luciferase assay. On the right, western blot confirmed expression of the transfected fusion protein, with HSP90 and actin as loading control.

**Fig. S2. Validation of DAB2IP HiBiT-tagged cells for high-throughput screening.**

**A** Workflow of CRISPR/Cas9-mediated gene knock-in. See material and methods for details. **B** Bulk HiBiT nucleofected cells are positive to luminescence. An identical number of the indicated cells were lysed and processed for luciferase assay. **C.** Bulk nucleofected cells display edited target sequence. Genomic DNA was amplified using primers designed to produce ca. 350 bp amplicons with a clear upshift in case of HiBiT integration. Parental genomic DNAs were amplified as control. **D.** DAB2IP knockdown reduces DAB2IP-HiBiT luciferase activity. Bulk PC3-HiBiT cells were transfected with DAB2IP siRNA or with DAB2IP-targeting miR-149-3p for 48 hours before luminescence reading. Data are mean ± SD of 3 wells per condition. On the right, western blot confirmed downregulation of DAB2IP protein. Actin was blotted as a loading control. **E.** Fluorescence intensity and luminescence intensity (DAB2IP levels) correlate with the number of cells. The indicated numbers of parental non-edited PC3 and PC3-HiBiT cells were plated in 384 well-plates. 24 hours later, cells were incubated with the CellTiterFluor (Promega) reagent and fluorescence was detected (left panel). After fluorescence detection, the same cells were incubated with NanoGlo lytic reagent (Promega) and luminescence was measured (right panel). Data are mean ± SD of three wells per condition (two-way ANOVA with Sidak correction). **F-G.** The Lum/Fluo ratio readily detects changes in DAB2IP protein levels. PC3-HiBiT **(F)** and DU145-HiBiT **(G)** cells were reverse transfected with DAB2IP siRNA in 384 well-plates. 72 hours after transfection, fluorescence and luminescence were read. Results are mean ± SD of 3 wells per condition (one-way ANOVA with Dunnett's post hoc).

**Fig. S3 Validation of identified drugs in DU145 prostate cancer cell line.**

**A** Histogram summarizes the Lum/Fluo ratio measured in DU145-HiBiT cells treated as in Fig. 1C. Data are mean ± SD of three indipendent experiments (one-way ANOVA with Dunnett's post hoc). **B** Dose-response curves of the three best candidate drugs in DU145-HiBiT cells. The graphs summarize the effect of each drug on DAB2IP levels (Lum/Fluo ratio) and the effect on viability (Fluorescence). The half-maximal effective concentration ($EC_{50}$) and half-maximal inhibitory concentration ($IC_{50}$) are also indicated. Data are mean of *n* independent experiments as indicated in figure ± SD. **C-D.** Effects of candidate drugs on DAB2IP in non-edited parental DU145. Cells were treated with drugs or DMSO (Ctrl) for 36h. When not indicated, drug concentration was 10uM. C. DAB2IP was detected by immunoblotting, with HSP90 as loading control. D. DAB2IP mRNA levels were measured by RT-qPCR in DU145 treated with the drugs for 36 hours. Values were normalized on histone H3 and compared to DMSO-treated controls. Data are mean ± SD of three independent experiments (**p* < 0.1, ns = non-significant; one-way ANOVA with Dunnett's post hoc).

**Fig. S4 The identified drugs likely act on DAB2IP via different molecular mechanisms.**

**A.** Perturbagens and Compounds significantly associated to Thio, Ube, or Sal in the Broad Institute ConnectivityMap accessed via the CLUE platform. **B.** Venn diagram summarizing the top 1500 genes affected by 24 hours treatment with 10 μM Thio, Ube, or Sal in MCF7 cells (from ConnectivityMap). **C.** For each drug, the top 1500 genes were used for gene set enrichment analysis (KEGG pathways) using the ShinyGO platform.

**Fig. S5 Hallmarks enriched in PC3 and MCF7 after drug treatment**.

**A** The top 1500 genes affected by 24 hours treatment with 10 μM Thio, Ube, or Sal in PC3 or MCF7 cells were used for gene set enrichment analysis (Hallmarks.MsigDB) using the ShinyGO platform.

#### Figure S1

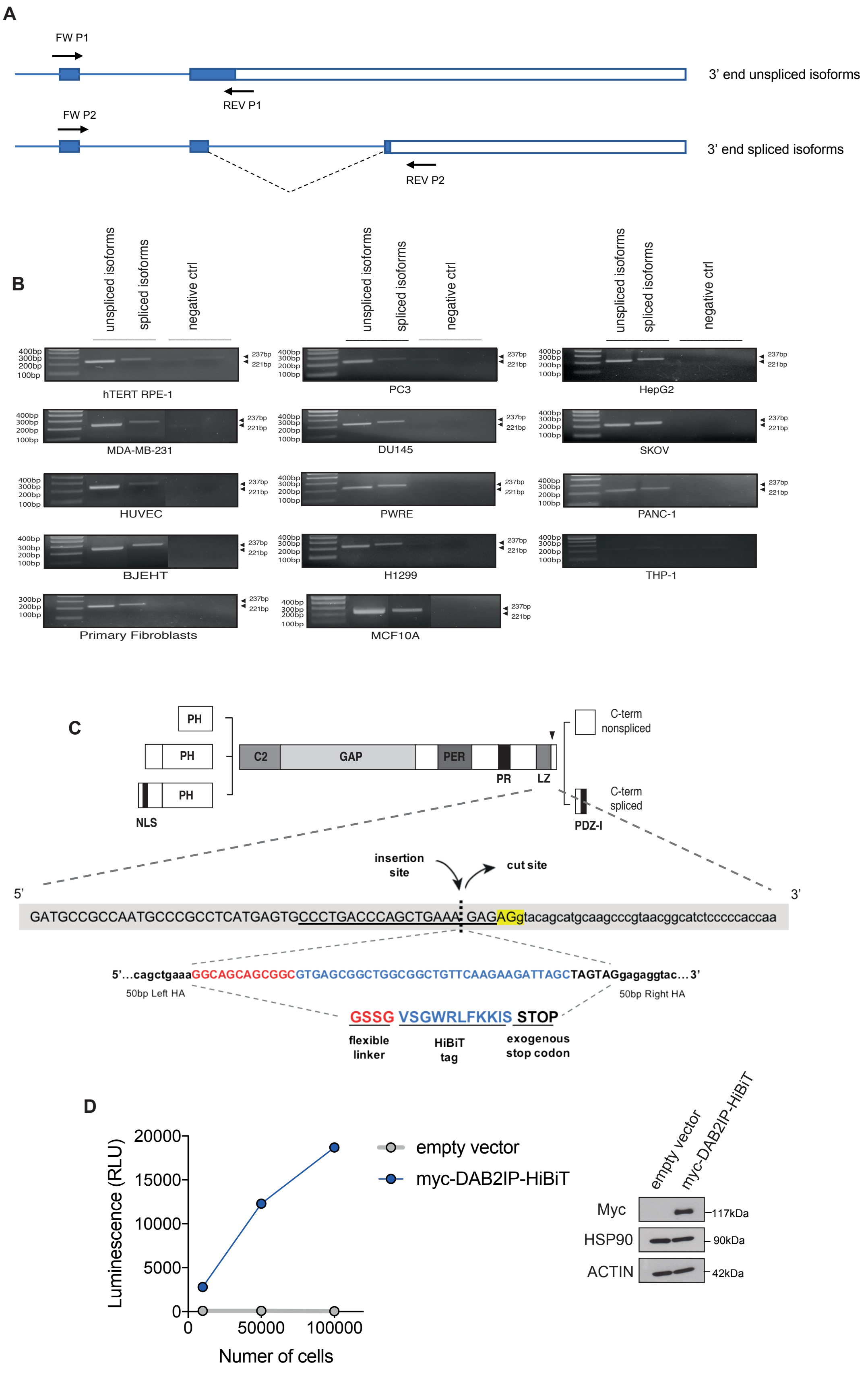
A
FW P1
3' end unspliced isoforms
REV P1
FW P2
3' end spliced isoforms
REV P2
B
unspliced isoforms
spliced isoforms
negative ctrl
400bp
300bp
200bp
100bp
237bp
221bp
hTERT RPE-1
PC3
HepG2
MDA-MB-231
DU145
SKOV
HUVEC
PWRE
PANC-1
BJEHT
H1299
THP-1
Primary Fibroblasts
MCF10A
C
PH
C2
GAP
PER
PR
LZ
NLS
C-term nonspliced
C-term spliced
PDZ-I
insertion site
cut site
5'
3'
GATGCCGCCAATGCCCGCCTCATGAGTGCCCTGACCCAGCTGAAA GAGAGgtacagcatgcaagcccgtaacggcatctcccccaccaa
5'...cagctgaaaGGCAGCAGCGGCGTGAGCGGCTGGCGGCTGTTCAAGAAGATTAGCTAGTAGgagaggtac...3'
50bp Left HA
50bp Right HA
GSSG
VSGWRLFKKIS
STOP
flexible linker
HiBiT tag
exogenous stop codon
D
Luminescence (RLU)
20000
15000
10000
5000
0
0
50000
100000
Numer of cells
empty vector
myc-DAB2IP-HiBiT
Myc
HSP90
ACTIN
117kDa
90kDa
42kDa

### Figure S2

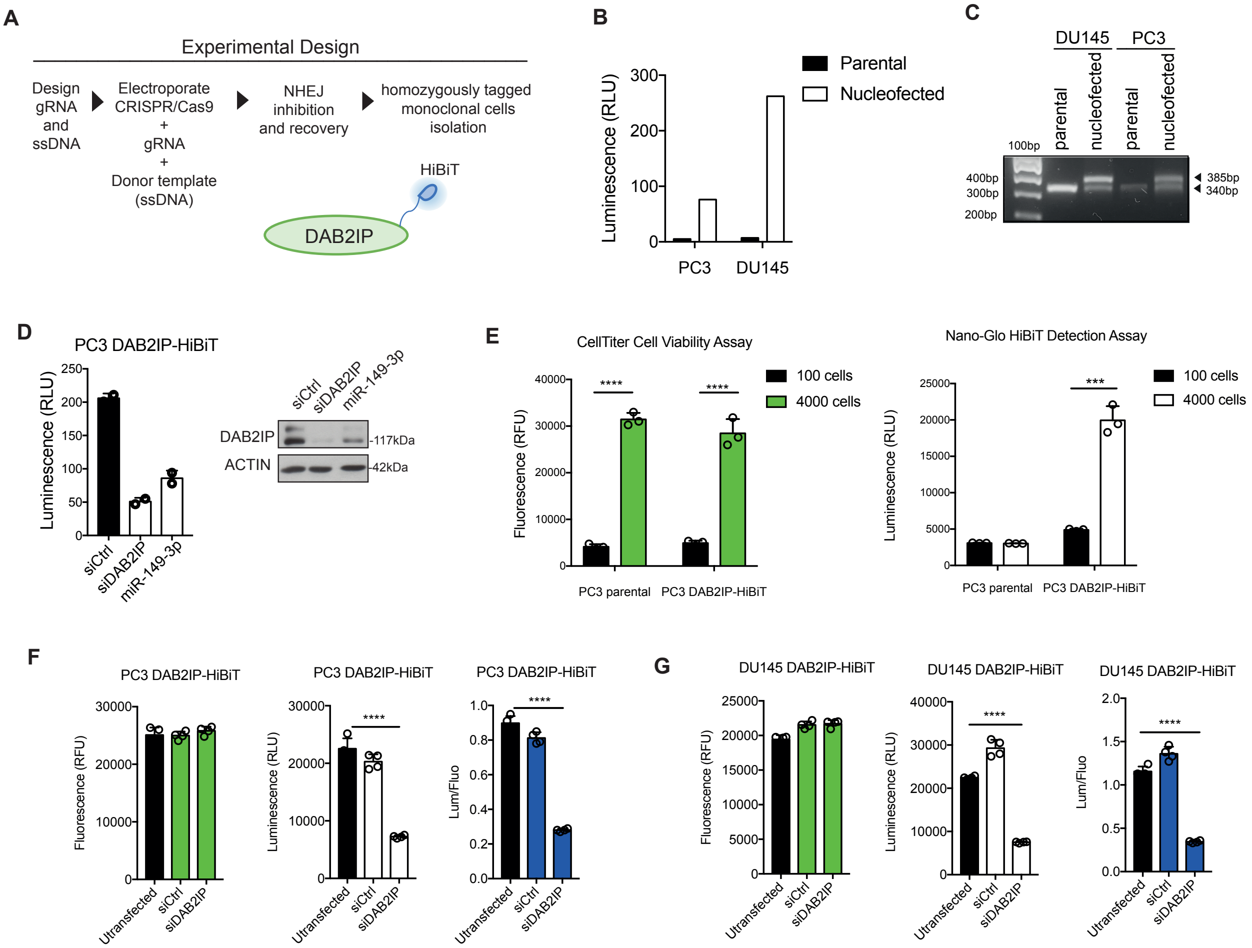
A
Experimental Design
Design gRNA and ssDNA
Electroporate CRISPR/Cas9 + gRNA + Donor template (ssDNA)
NHEJ inhibition and recovery
homozygously tagged monoclonal cells isolation
HiBiT
DAB2IP
B
Parental
Nucleofected
Luminescence (RLU)
PC3
DU145
C
DU145
PC3
100bp
parental
nucleofected
parental
nucleofected
400bp
300bp
200bp
385bp
340bp
D
PC3 DAB2IP-HiBiT
Luminescence (RLU)
siCtrl
siDAB2IP
miR-149-3p
DAB2IP
ACTIN
117kDa
42kDa
E
CellTiter Cell Viability Assay
100 cells
4000 cells
Fluorescence (RFU)
PC3 parental
PC3 DAB2IP-HiBiT
Nano-Glo HiBiT Detection Assay
Luminescence (RLU)
F
PC3 DAB2IP-HiBiT
Fluorescence (RFU)
Luminescence (RLU)
Lum/Fluo
Utransfected
siCtrl
siDAB2IP
G
DU145 DAB2IP-HiBiT

### Figure S3

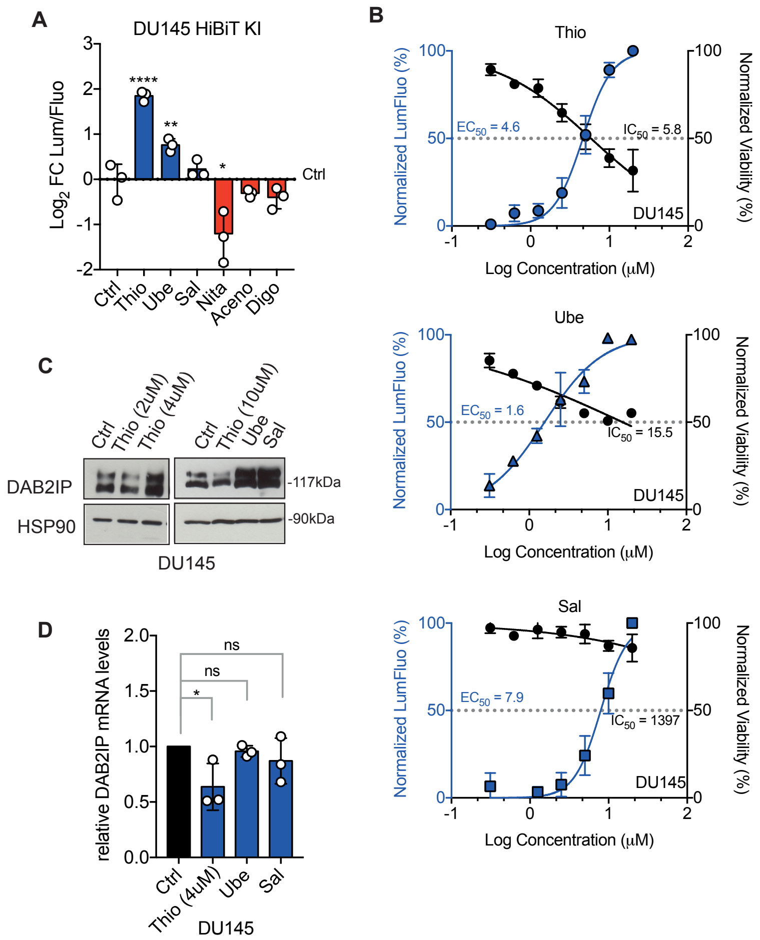

A
DU145 HiBiT KI
Log2 FC Lum/Fluo
****
**
*
Ctrl
Ctrl
Thio
Ube
Sal
Nita
Aceno
Digo
B
Thio
Normalized LumFluo (%)
Normalized Viability (%)
EC50 = 4.6
IC50 = 5.8
DU145
Log Concentration (µM)
Ube
EC50 = 1.6
IC50 = 15.5
DU145
Log Concentration (µM)
Sal
EC50 = 7.9
IC50 = 1397
DU145
Log Concentration (µM)
C
Ctrl
Thio (2uM)
Thio (4uM)
Ctrl
Thio (10uM)
Ube
Sal
DAB2IP
HSP90
-117kDa
-90kDa
DU145
D
relative DAB2IP mRNA levels
ns
ns
*
Ctrl
Thio (4uM)
Ube
Sal
DU145

### Figure S4

**A**

#### Thiostrepton

**Perturbagen class**

| score | type | name |
|---|---|---|
| 99.89 | | IKK inhibitor |
| 99.81 | | JAK inhibitor |
| 99.70 | | FLT3 inhibitor |
| 99.56 | | PKA inhibitor |
| 99.52 | | HMGCR inhibitor |
| -15.74 | | PI3K Signaling LOF |
| -16.91 | | Bile acid |
| -31.48 | | NADH ubiquinone oxidoreductase supernumerary subunits LOF |
| -42.09 | | Phospholipases LOF |
| -90.71 | | Integrin subunits beta LOF |

**Compound**

| score | type | name |
|---|---|---|
| 99.97 | | thiostrepton |
| 99.37 | | arachidonyl-trifluoro-methane |
| 99.33 | | BX-795 |
| 99.26 | | JAK3-inhibitor-VI |
| 99.12 | | IKK-16 |
| 99.10 | | NSC-632839 |
| 98.98 | | cucurbitacin-i |
| 98.94 | | BX-912 |
| 98.94 | | purvalanol-a |
| 98.91 | | diphencyprone |
| -61.60 | | PD-169316 |
| -64.94 | | PP-30 |
| -64.96 | | SB-525334 |
| -67.28 | | AGK-2 |
| -68.22 | | PF-04217903 |
| -69.14 | | BAY-K8644 |
| -70.17 | | evodiamine |
| -73.36 | | NF-449 |
| -80.17 | | navitoclax |
| -86.88 | | dictamnine |

#### Ubenimex

**Perturbagen class**

| score | type | name |
|---|---|---|
| 99.28 | | Rho associated kinase inhibitor |
| 98.62 | | Sigma receptor antagonist |
| 97.01 | | Sterol demethylase inhibitor |
| 95.47 | | Estrogen receptor agonist |
| 95.25 | | Tachykinin antagonist |
| 3.80 | | Sirtuins GOF |
| 3.36 | | PRD class homeoboxes and psudogenes LOF |
| -15.43 | | NADH ubiquinone oxidoreductase supernumerary subunits LOF |
| -33.80 | | Phospholipases LOF |
| -40.93 | | Integrin subunits beta LOF |

**Compound**

| score | type | name |
|---|---|---|
| 99.99 | | ubenimex |
| 99.93 | | BRL-15572 |
| 99.93 | | ginsenoside |
| 99.93 | | rifampicin |
| 99.93 | | naproxen |
| 99.93 | | EBPC |
| 99.93 | | bethanechol |
| 99.93 | | JAK3-inhibitor-VI |
| 99.93 | | aminogenistein |
| 99.93 | | arachidonamide |
| -54.10 | | danoprevir |
| -58.25 | | QW-BI-011 |
| -60.66 | | tyrphostin-AG-1295 |
| -67.40 | | tranilast |
| -67.75 | | sitagliptin |
| -77.18 | | bendroflumethiazide |
| -78.67 | | norepinephrine |
| -79.21 | | griseofulvin |
| -95.89 | | ofloxacin |
| -99.82 | | BRD-A80383043 |

#### Salmeterol

**Perturbagen class**

| score | type | name |
|---|---|---|
| 98.88 | | HMGCR inhibitor |
| 98.71 | | Beta-adrenergic receptor agonist |
| 98.52 | | Progesterone receptor agonist |
| 98.27 | | Aromatase inhibitor |
| 97.98 | | Tricyclic antidepressant |
| 3.38 | | Estrogen receptor agonist |
| 3.24 | | Retinoid receptor agonist |
| 2.51 | | Non Homologous End Joining LOF |
| 2.09 | | MCM family LOF |
| 1.64 | | Supressors of cytokine signaling GOF |

**Compound**

| score | type | name |
|---|---|---|
| 99.99 | | salmeterol |
| 99.68 | | sertraline |
| 99.65 | | latanoprost |
| 99.58 | | amlodipine |
| 99.12 | | cerivastatin |
| 98.80 | | metergoline |
| 98.80 | | heliomycin |
| 98.57 | | chlormadinone |
| 97.99 | | pimozide |
| 97.53 | | forskolin |
| -30.18 | | trapidil |
| -33.97 | | cosmosiin |
| -37.91 | | RHO-kinase-inhibitor-III[rockout] |
| -41.78 | | ziprasidone |
| -44.05 | | CDC |
| -53.25 | | JNJ-10191584 |
| -57.74 | | fluticasone |
| -60.28 | | zaprinast |
| -67.29 | | clofibric-acid |
| -83.96 | | isoflupredone |

**B**

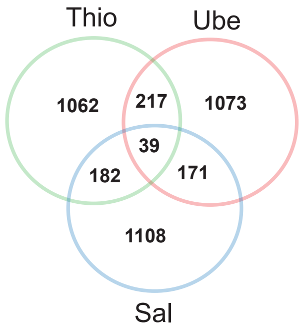


**C**

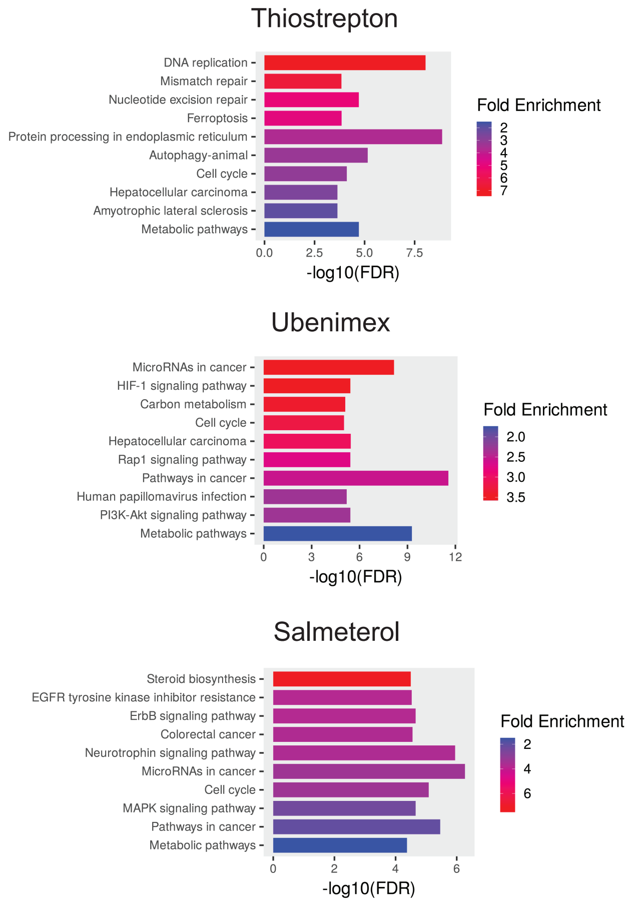

### Figure S5

A

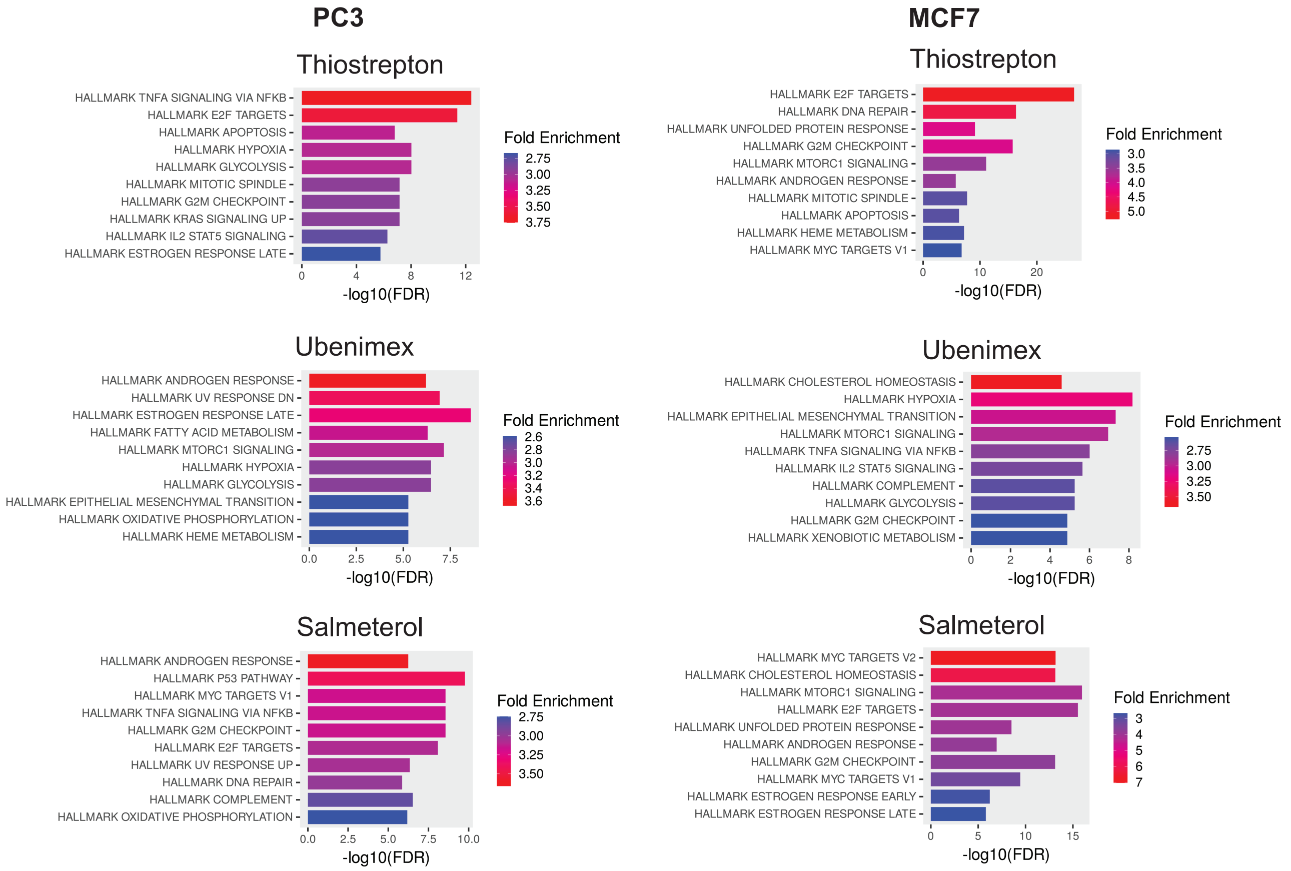
PC3
MCF7
Thiostrepton
HALLMARK TNFA SIGNALING VIA NFKB
HALLMARK E2F TARGETS
HALLMARK APOPTOSIS
HALLMARK HYPOXIA
HALLMARK GLYCOLYSIS
HALLMARK MITOTIC SPINDLE
HALLMARK G2M CHECKPOINT
HALLMARK KRAS SIGNALING UP
HALLMARK IL2 STAT5 SIGNALING
HALLMARK ESTROGEN RESPONSE LATE
-log10(FDR)
Fold Enrichment
Thiostrepton
HALLMARK E2F TARGETS
HALLMARK DNA REPAIR
HALLMARK UNFOLDED PROTEIN RESPONSE
HALLMARK G2M CHECKPOINT
HALLMARK MTORC1 SIGNALING
HALLMARK ANDROGEN RESPONSE
HALLMARK MITOTIC SPINDLE
HALLMARK APOPTOSIS
HALLMARK HEME METABOLISM
HALLMARK MYC TARGETS V1
-log10(FDR)
Fold Enrichment
Ubenimex
HALLMARK ANDROGEN RESPONSE
HALLMARK UV RESPONSE DN
HALLMARK ESTROGEN RESPONSE LATE
HALLMARK FATTY ACID METABOLISM
HALLMARK MTORC1 SIGNALING
HALLMARK HYPOXIA
HALLMARK GLYCOLYSIS
HALLMARK EPITHELIAL MESENCHYMAL TRANSITION
HALLMARK OXIDATIVE PHOSPHORYLATION
HALLMARK HEME METABOLISM
-log10(FDR)
Fold Enrichment
Ubenimex
HALLMARK CHOLESTEROL HOMEOSTASIS
HALLMARK HYPOXIA
HALLMARK EPITHELIAL MESENCHYMAL TRANSITION
HALLMARK MTORC1 SIGNALING
HALLMARK TNFA SIGNALING VIA NFKB
HALLMARK IL2 STAT5 SIGNALING
HALLMARK COMPLEMENT
HALLMARK GLYCOLYSIS
HALLMARK G2M CHECKPOINT
HALLMARK XENOBIOTIC METABOLISM
-log10(FDR)
Fold Enrichment
Salmeterol
HALLMARK ANDROGEN RESPONSE
HALLMARK P53 PATHWAY
HALLMARK MYC TARGETS V1
HALLMARK TNFA SIGNALING VIA NFKB
HALLMARK G2M CHECKPOINT
HALLMARK E2F TARGETS
HALLMARK UV RESPONSE UP
HALLMARK DNA REPAIR
HALLMARK COMPLEMENT
HALLMARK OXIDATIVE PHOSPHORYLATION
-log10(FDR)
Fold Enrichment
Salmeterol
HALLMARK MYC TARGETS V2
HALLMARK CHOLESTEROL HOMEOSTASIS
HALLMARK MTORC1 SIGNALING
HALLMARK E2F TARGETS
HALLMARK UNFOLDED PROTEIN RESPONSE
HALLMARK ANDROGEN RESPONSE
HALLMARK G2M CHECKPOINT
HALLMARK MYC TARGETS V1
HALLMARK ESTROGEN RESPONSE EARLY
HALLMARK ESTROGEN RESPONSE LATE
-log10(FDR)
Fold Enrichment
